# Neural responses to biological motion distinguish autistic and schizotypal traits in the general population

**DOI:** 10.1101/2022.03.24.485704

**Authors:** Matthew Hudson, Severi Santavirta, Vesa Putkinen, Kerttu Seppälä, Lihua Sun, Tomi Karjalainen, Henry K. Karlsson, Jussi Hirvonen, Lauri Nummenmaa

**Author notes:** **Declaration of interests:** The authors declare no competing interests. **Funding:** The research was funded by grants awarded to LN from the Academy of Finland (#294897 and #332225) and a European Research Council Starting Grant (#313000). Corresponding Author: School of Psychology, University of Plymouth, Plymouth, Devon, UK, PL4 8AA.

## Abstract

Difficulties in social interactions are common to both autism and schizophrenia, and contribute to correlated autistic and schizotypal traits in the neurotypical population. It remains unresolved whether this represents a shared etiology or a superficial phenotypic overlap. Both conditions are associated with atypical neural activity in response to the perception of social stimuli, and also decreased neural synchronization between individuals that may prohibit establishing shared experiences. This study sought to establish if neural activity and neural synchronization associated with biological motion perception are differentially associated with autistic and schizotypal traits in the neurotypical population. Participants watched an audiovisual montage of naturalistic social interactions whilst hemodynamic brain activity was measured with fMRI. A separate sample of participants provided a continuous measure of the extent of biological motion, which was used to predict hemodynamic activity. General Linear Model analysis revealed that biological motion perception was associated with neural activity across the action-observation network. However, inter-subject phase synchronization analysis revealed that neural activity synchronized between individuals in occipital and parietal areas, but de-synchronized in temporal and frontal regions. Autistic traits were associated with a decrease in neural activity (precuneus, middle cingulate gyrus) and schizotypal traits were associated with a decrease in neural synchronization (middle and inferior frontal gyri). Biological motion perception elicits convergent and divergent patterns of neural activity and neural synchronization, and are differentially associated with shared traits related with autism and schizophrenia in the general population, suggesting that they originate from different neural mechanisms.

---

Autism and schizophrenia are both characterized by difficulties in social interactions (American Psychiatric Association, 2013) as well as atypical perceptual and cognitive processes supporting social cognition (Abdi & Sharma, 2014; Hommer & Swedo, 2015; King & Lord, 2011; Pina-Camacho, Parellada, & Kyriakopoulos, 2016; Sasson, Pinkham, Carpenter, & Belger, 2011). Whilst these conditions can be readily distinguished by the other symptom clusters (such as restricted interests and behaviors in autism, and delusions and hallucinations in psychosis), the phenotypic convergence of social features contributes to diagnostic uncertainty (Nylander, 2014), mutual comorbidity (Barneveld et al., 2011; De Crescenzo, et al., 2017; Kincaid, Doris, Shannon, & Mulholland, 2017; Kiyono, et al., 2020; Lugo-Marína et al., 2019; Zheng, Zheng, & Zou, 2018), and heritability (Sullivan, et al., 2012; Wieckowski, Mukhtar, Lee, Xing, & Walker, 2017). The degree of autistic and schizotypal traits varies in the general population (Landry & Chouinard, 2016; van Os & Reininghaus, 2016), and these dimensions are also correlated, especially with respect to social behavior (Zhou, et al., 2019). However, it is unclear whether this convergence reflects a shared etiology or dissociable etiologies that manifest in overlapping phenotypes (Chisholm, Linb, Abu-Akela, & Wood, 2015; DeVylder & Oh, 2014). A key issue is therefore to identify neurocognitive correlates of social cognition and perception that may discriminate between them.

## Social cognition in autism and schizophrenia

Social cognition encompasses the learning, communicative, motivational, identification, and predictive processes that characterize our interpersonal relationships, and involves distributed processes that work in concert to integrate this complex information dynamically along timespans from milliseconds to years (Molapour et al., 2021). These complex behaviors require input from more fundamental perceptual processes (Pitcher & Ungerleider, 2021; Ramsey & Ward, 2020). The distinctive kinematic profile of biological motion, and the social cues inherent in facial and bodily movements, allow one to not only detect intentional agents, but to spontaneously and implicitly infer the hidden mental states that are driving their behavior. Difficulties in interpreting the agency, intentionality, emotion, and purpose of other’s behavior (Couture et al., 2010; Pinkham et al., 2019) may be the precipitating feature that makes complex social reasoning even more difficult in autism and schizophrenia (Froese, Stanghellini, & Bertelli, 2013; Gallagher & Varga, 2015). Indeed, people with either condition are able to detect biological motion and discriminate different actions, such as dancing vs fighting (Cusack, Williams, & Neri, 2015) and walking direction (Keane, Peng, Demmin, Silverstein, & Lu, 2018), but show a difficulty in making higher-level inferences regarding the emotional and intentional dispositions of actors depicted (Corrigan, 1997; Hudson, Burnett, & Jellema, 2012; Hudson, Nicholson, Kharko, McKenzie, & Bach, 2021; Kaiser & Shiffrar, 2009; Okruszek & Pilecka, 2017; Savla, Vella, Armstrong, Penn, & Twamley, 2013; Todorova, McBean Hatton, & Pollick, 2019). Moreover, these abilities correlate with autistic and schizotypal traits in the general population (Blain, Peterman, & Park, 2017; Gray, Jenkins, Heberlein, & Wegner, 2011; Hudson, Nijboer, & Jellema, 2012). Social perceptual processes may therefore provide the origin for the high-level social difficulties evident in autism and schizophrenia, and potentially present a point of divergence between the neurophenotypes of the two conditions.

## Neural mechanisms of social perception in Autism and Schizophrenia

In neurotypical individuals, the visual perception of social stimuli reliably elicits widespread neural activity in a distributed and hierarchical network of cortical regions, from category specific regions of early visual areas, to motion processing regions sensitive to the intentionality of others actions, and visuomotor areas of the parietal and prefrontal cortices (Caspers, Zillesa, Laird, & Eickhoff, 2010; Decety & Grèzes, 1999; Grosbras, Beaton, & Eickhoff, 2012; Pavlova, 2012). The neural activity is linearly related with the frequency and prominence of facial/bodily motion that can be seen (Bartels & Zeki, 2004; Lahnakoski et al., 2012). Moreover, neural activity becomes synchronized between individuals in many of these areas during the perception of complex social interactions (Bolt, Nomi, Vij, Chang, & Uddin, 2018; Hasson, Nir, Levy, Fuhrmann, & Malach, 2004). Neural synchronization provides a measure of how voxel-specific neural activity is associated with complex and dynamic multidimensional stimuli that are representative of real-world experience, especially those encountered in social interactions (Adolphs, Nummenmaa, Todorov, & Haxby, 2016; Hasson & Frith, 2016; Zaki & Ochsner, 2009). Such synchronization may reflect, at least, a population-wide reliability in the neural response to a common stimulus feature (Hudson et al., 2020), and at most, may provide the basis by which we share an understanding of the world with other people that is necessary for many key high-level social functions (Nummenmaa, Lahnakoski, & Glerean, 2018; Redcay & Moraczewskib, 2020). However, it is unknown whether the synchronization of neural activity correlates with the extent and intensity of biological motion being observed, and how this may differ from the magnitude of activity typically assessed as a neural correlate of social perception.

Differential patterns of neural magnitude and synchronization in response to social stimuli may provide important insights into individual differences in social behaviours that are associated with autism and schizophrenia. Those with autism and schizophrenia consistently exhibit atypical neural activity in the network of regions implicated in social perception (Chan & Han, 2020; Glerean et al., 2016; Jáni & Kašpárek, 2017; Mehta, Thirthalli, Aneelraj, Jadhav, Gangadhar, & Keshavan, 2014; Philip, Dauvermann, Whalley, Baynham, Lawrie, & Stanfield, 2012; Yang & Hofmann, 2016). Direct comparisons between those with autism and schizophrenia have revealed comparably atypical neural activity when perceiving a range of social stimuli (Abdi & Sharma, 2014; King & Lord, 2011; Sasson, Pinkham, Carpenter, & Belger, 2011). However, a trend for hyperactivity in schizophrenia, and hypoactivity in autism, as well as hyper and hypo connectivity within these networks respectively, hints at dissociable neural mechanisms underpinning very similar behavioral features (Barlati, Minelli, Ceraso, Nibbio, Silva, Deste, Turrina, & Vita, 2020; Ciaramidaro, Bölte, Schlitt, Hainz, Poustka, Weber, Bara, Freitag, & Walter, 2015; Eack, Wojtalik, Keshavan, & Minshew, 2017; Sugranyes, Kyriakopoulos, Corrigall, Taylor, & Frangou, 2011). However, neural activity associated with biological motion perception is positively correlated with both autistic (Puglia & Morris, 2017; Thurman, van Boxtel, Monti, Chiang, & Lu, 2016) and schizotypal traits (Hur, Blake, Cho, Kim, Kim, Choi, Kang, & Kwon, 2016; Platek, Fonteyn, Izzetoglu, Myers, Ayaz, Li, & Chance, 2005), arguing for a dissociation between typical and atypical levels of these traits in the neural mechanisms underpinning perceptual processes of social cognition.

Moreover, there is a decreased neural synchronization with other people in both autism (Hasson, Avidan, Gelbard, Vallines, Harel, Minshew, & Behrmann, 2009; Salmi et al., 2009) and schizophrenia (Lerner et al., 2018; Mäntylä, Nummenmaa, Rikandi, Lindgren, Kieseppä, Hari, Suvisaari, & Raij, 2018) that may imply a more idiosyncratic and variable perception and interpretation of the world, and which may contribute to a difficulty in establishing a shared state of mind with other people. However, no studies have assessed neural synchronization in response to the perception of social stimuli and how this is related to atypical social behavior associated with autism and schizophrenia.

## The Current Study

The aims of this study are twofold. Firstly, to establish how the extent of neural synchronization between individuals during the perception of biological motion converges or diverges from the magnitude of neural activity that traditionally defines the action observation network. Theoretically, the amplitude of the neural signal is independent of the between-subject synchronization, even if they are both associated with the same stimulus features (Nastase, Gazzola, Hasson, & Keysers, 2019). Furthermore, measures of neural synchrony can reveal regions of neural response that GLM approaches do not (Hejnar, Kiehl, & Calhoun, 2007; Pajula, Kauppi, & Tohka, 2012; Xu, Bolt, Nomi, Li, Zheng, Fu, Kendrick, Becker, & Uddin, 2020). There is therefore a compelling reason to establish how both the amplitude and reliability of neural activity varies with biological motion perception. To this end, we had participants view a series of movie clips of complex and dynamic naturalistic social interactions in an fMRI scanner. These clips were observed by a separate sample of participants who gave a continuous rating of the extent of biological motion present, which provided a behavioral measure that was correlated with both the magnitude and synchronization of neural activity in each voxel.

The second aim was to investigate how differential patterns of neural magnitude and synchronization associated with biological motion perception may differentiate the overlapping social difficulties characterized by autistic and schizotypal traits in the general population. Each participant completed the Autistic Spectrum Quotient (AQ: Baron-Cohen, Wheelwright, Skinner, Martin, & Clubley, 2001) and Oxford-Liverpool Inventory of Feelings and Experiences (O-LIFE: Mason, Linney, & Claridge, 2005) to measure autistic and schizotypal traits respectively. We were specifically interested not only in the overall scores, but the sub-scales indicative of social behavior – the social skills sub-scale of the AQ and the introversion sub-scale of the O-LIFE – and the extent to which they were associated with individual differences in the magnitude of neural activity and neural synchronization in response to biological motion perception.

In doing so, we aim to provide a detailed account of the neural mechanisms of core social perception processes that encompasses not only how neural activity is associated with biological motion perception per se, but how it is associated with the neural activity of other people watching the same social interaction. Moreover, we aim to show how these potentially convergent or divergent patterns of neural activity may differentiate individual differences in social behavior associated with autistic and schizotypal traits in the general population, and provide insight into how the overlapping atypical social abilities of autism and schizophrenia may be discerned at the neural level.

## Method

### Participants

104 participants took part in the study and were recruited from the University of Turku and wider community. Two were excluded due to scanning artefacts, two were excluded due to gross brain abnormalities, and three were excluded due to incomplete questionnaire data, leaving 97 participants in the analysis (48 females, age mean = 31.3 years, SD = 9.4). All participants were screened with standard MRI exclusion criteria, in addition to neurological, neuropsychiatric, and psychotropic substance use related contraindications. Participants gave written informed consent prior to the study and were paid for participation. The study was approved by the ethics committee of the Hospital District of South West Finland in accordance with the declaration of Helsinki.

## Materials and Stimuli

### Questionnaires

Participants completed the questionnaires online after the scanning session. The AQ (Baron-Cohen, Wheelwright, Skinner, Martin, & Clubley, 2001) is a 50 item self-report questionnaire that measures autistic traits in the general population, and contains five sub-scales relating to social skills, attention to detail, attention switching, imagination, and communication. Responses are scored as 1 or 0, with higher scores indicating greater autistic traits, with a maximum score of 50. The O-LIFE (Mason, Linney, & Claridge, 2005) is a 43 item self-report questionnaire that measures schizotypal traits in the general population, and contains four sub-scales relating to introvertive anhedonia, cognitive disorganization, unusual experiences, and impulsive non-conformity. Responses are scored as 1 or 0, with higher scores indicating greater schizotypal traits, with a maximum score of 43. The Finnish language translations showed comparably high internal consistency (Cronbach’s alpha coefficient: AQ = .711; OLIFE = .842) to previous English language versions (Baron-Cohen, Wheelwright, Skinner, Martin, & Clubley, 2001; Mason, Linney, & Claridge, 2005). Previous research has shown that the AQ and OLIFE correlate positively in the general population, especially with respect to the social skills and introvertive anhedonia sub-scales that measure social behavior (Russell-Smith, Bayliss, & Maybery, 2011).

### Biological Motion Stimulus

During the fMRI scan the participants watched an audio-visual montage of 96 clips taken from popular movies with English speech (mean clip duration 11.5 secs, total duration 19.6 mins); the participants had no specific task other than to pay attention to the movie. The montage was designed to provide a high dimensional representation of complex naturalistic social and emotional interactions, and has been validated in previous studies (Lahnakoski et al. 2012; Karjalainen et al. 2017, 2019). The clips were presented in a fixed order for each participant to allow for the inter-subject synchronization analysis. A fixation cross was presented at the start (5.2 secs) and end (15.6 secs) of the run. Stimulus video was displayed using goggles affixed to the head coil (NordicNeuroLab VisualSystem). Audio was played through SensiMetrics S14 earphones (100 Hz–8 kHz bandwidth, 110 dB SPL). Volume was adjusted individually to a comfortable level that could still be heard over the scanner noise.

The montage was viewed by five separate participants who provided a continuous rating for the presence of biological motion (Figure 1). The stimulus was presented on a computer monitor and headphones, whilst the participant provided ratings by moving the mouse forward for increased presence or backward for decreased presence. Ratings were taken at 0.25Hz intervals and down-sampled to match the TR of the fMRI time series to be used as a regressor of interest to establish the relationship between biological motion perception and the magnitude and synchronization of neural activity. Inter-class correlation analysis (*r* = .57) indicated good reliability between raters.

**Figure 1.**
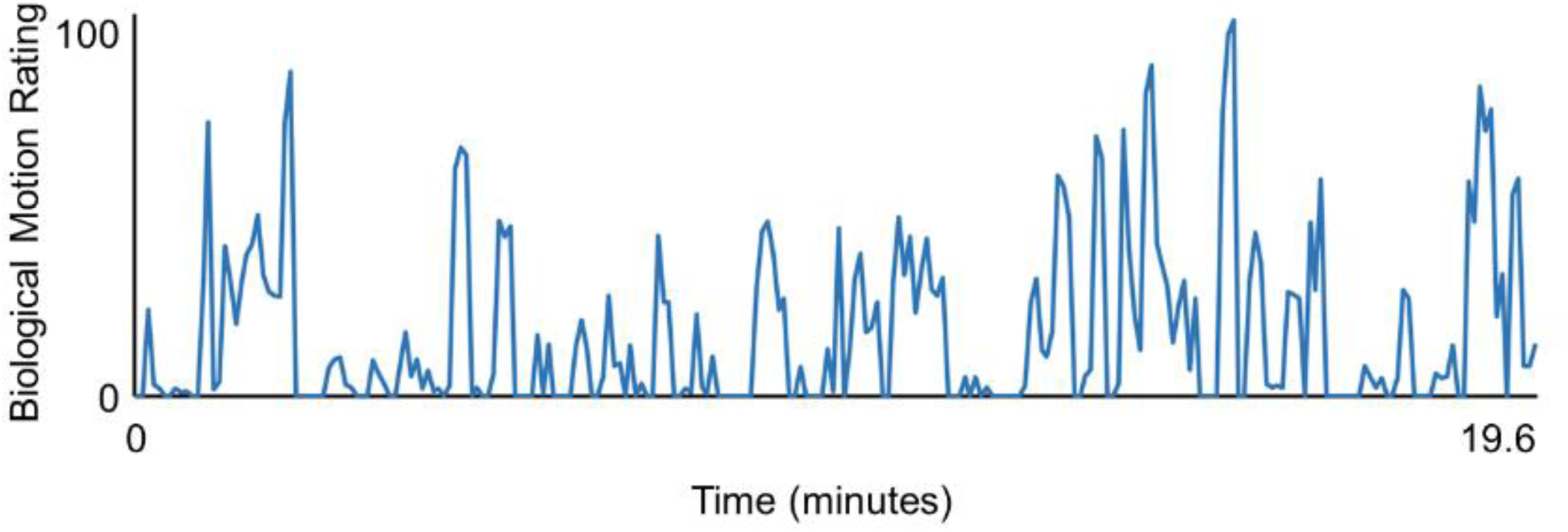
Average viewer ratings of the extent of biological motion in the stimulus.

#### Procedure

### MRI Data Acquisition and Preprocessing

MRI scanning took place at Turku PET Centre at the University of Turku using a Phillips Ingenuity TF PET/MR 3-T whole-body scanner. High-resolution (1 mm3) structural images were obtained with a T1-weighted sequence (TR 9.8 ms, TE 4.6 ms, flip angle 7°, 250 mm FOV, 256 × 256 reconstruction matrix). A total of 467 functional volumes were acquired, with a T2∗-weighted echo-planar imaging sequence (TR 2600 ms, TE 30 ms, 75° flip angle, 240 mm FOV, 80 × 80 reconstruction matrix, 62.5 kHz bandwidth, 3.0 mm slice thickness, 45 interleaved slices acquired in ascending order without gaps).

Preprocessing of MRI data used fMRIPprep 1.3.0.2 (Esteban et al., 2019). The anatomical T1-weighted reference image was subject to correction for intensity non-uniformity, skull stripping, brain surface reconstruction, and spatial normalization to the ICBM 152 Nonlinear Asymmetrical template version 2009c (Fonov et al., 2009) using nonlinear registration with antsRegistration (ANTs 2.2.0) and brain tissue segmentation. The function data were subject to coregistration to the T1w reference, slice-time correction, spatial smoothing with a 6-mm Gaussian kernel, automatic removal of motion artifacts using ICA-AROMA (Pruim et al. 2015), and resampling of the MNI152NLin2009cAsym standard space. Low-frequency drifts were removed with a 240-s-Savitzky–Golay filter (Çukur et al. 2013).

### Data Analysis

We conducted both General Linear Model (GLM) and Inter-Subject Phase Synchronization (ISPS) analyses to establish how biological motion perception is associated with regionally specific changes in BOLD activity and neural synchronization between individuals respectively. In addition, we repeated these analyses with questionnaire scores as participant level regressors to establish how BOLD activity and neural synchronization between individuals associated with biological motion perception varies with autistic and schizotypal traits. We also sought to establish how functional connectivity during biological motion perception is related to autistic and schizotypal traits. To this end, Seed-Based Phase Synchronization (SBPS) was employed, which takes the instantaneous synchronization of BOLD activity between region pairs for each participant and correlated this with the biological motion regressor and questionnaire scores.

Based on our hypotheses, we focused on two conjunctions of trait measures. Firstly, total AQ and OLIFE scores were entered as orthogonalized regressors in the GLM, ISPS, and SBPS analyses. Secondly, the social skills and introvertive subscales of the AQ and OLIFE (respectively) were entered as orthogonalized regressors. In each analysis, the two scores were entered as regressors, with each acting as a regressor of no-interest of the other. This enabled the investigation of the independent contributions of these traits to explain neural response to biological motion, despite high covariance between the scores themselves. Further exploratory analyses using sub-scale scores within each trait measure (e.g. all sub-scales of the AQ and all of the OLIFE), and inter-trait relationships between sub-scales revealed to be highly correlated (e.g., between Communication [AQ] and Impulsive Non-Conformity [OLIFE]) can be found in the Supplementary Materials.

### General Linear Model Analysis

GLM analyses were conducted with SPM12 (www.fil.ion.ucl.ac.uk/spm) with a two-stage random effects analysis. The biological motion ratings were convolved with a canonical hemodynamic response function and entered as a regressor into the first-level GLM analysis, using a high-pass filter of 128s. The results of each participant in the first-level analysis were entered into a second-level random effects analysis using a one-sample t-test, with a FWE alpha threshold of *p* < .001. The questionnaire scores were entered as a participant level regressor in the second-level analysis, with a cluster level FDR threshold after an uncorrected voxel threshold of *p* < .001.

### Inter-Subject Phase Synchronization Analysis

The data were preprocessed for phase synchronization analysis using the FunPsy toolbox (https://github.com/eglerean/funpsy) and described in detail in Glerean et al. (2012). For each participant, the voxel specific time series was band-pass filtered (0.04-0.07 Hz), and the phase analytic signal (in radians) of the Hilbert transformed BOLD response of each voxel was calculated. The phase analytic signal at each timepoint was subtracted (and inversed) from that of the equivalent voxel from each of the other participants, and then averaged, to produce a 4D (space X time) measure of phase similarity of each participant with the rest of the sample. As the phase similarity measure is instantaneous, it provides a more temporally precise indicator of neural synchronization than the sliding window analyses of inter-subject correlation.

Three ISPS analyses were conducted. (1) To establish how neural synchronization *per se* varies as a function of individual differences in autistic and schizotypal traits (i.e. irrespective of stimulus features) the total AQ and OLIFE scores were entered as participant level regressors, as were the social skills and introvertive sub-scales, with a cluster level threshold after an uncorrected voxel threshold of *p* < .001. (2) The relationship between ISPS and biological motion perception was investigated by taking each participants voxel specific phase similarity time series and correlating it with the HRF convolved biological motion regressor for each voxel. The resulting r values were fisher z transformed to make them suitable for statistical analysis. These were then entered into a second level analysis in SMP12 using a one-sample t-test and a FWE corrected alpha threshold of *p* < .001. (3) The questionnaire scores were entered as a participant level regressor to establish how neural synchronization (ISPS) associated with biological motion perception varies with autistic and schizotypal traits, with a cluster level threshold after an uncorrected voxel threshold of *p* < .001.

### Seed-Based Phase Synchronization

Regions of interest were defined according to the Brainnettome atlas (Fan et al., 2016), consisting of 123 cortical regions per hemisphere and 28 cerebellar regions (although the Cerebellum_Vermis_Crus_I could not be identified in our sample and was excluded, leaving 27 cerebellar regions). The BOLD signal in each region was averaged across voxels, and the band-pass filtered (0.04-0.07 Hz) time series was converted to a phase analytic signal. For each participant, the instantaneous phase similarity between each region pair was calculated to provide a continuous measure of neural synchronization between regions within each subject (region [273] X region [273] by time [467]).

Three SBPS analyses were conducted. (1) To establish how SBPS varies with biological motion perception, the phase similarity was averaged across participants at each timepoint to provide a sample wide measure of between-region synchronization at each time point. Each time series was then correlated with the HRF convolved biological motion regressor, with a corrected df that accounted for autocorrelations between the phase similarity and the biological motion regressor (Pyper and Peterman, 1998), with an FDR corrected *p* < .001. (2) The extent to which the association between biological motion perception and SBPS varies with individual differences in autistic and schizotypal traits was investigated by correlating the HRF convolved biological motion regressor with the phase similarity of each region pair for each participant individually. The Fischer z-transformed r values were then correlated across participants with the questionnaire scores at each node, with a FDR corrected *p* < .001. (3) Lastly, the association of autistic and schizotypal traits with SBPS, irrespective of stimulus features, was established by averaging the phase similarity of each region pair across the temporal dimension for each participant, to provide an overall measure of between-region synchronization across the scanning run. Each node was then correlated with the questionnaire scores with an FDR corrected *p* <.001.

### Data Availability

Thresholded and unthresholded results of the GLM analyses and inter-subject phase synchronization analyses are available on NeuroVault (https://identifiers.org/neurovault.collection:12349). Thresholded and unthresholded correlation matrices for the seed-based phase synchronization analyses, and questionnaire data for the AQ and OLIFE (totals and subscales), are available at https://osf.io/vg8zx/.

## Results

### Questionnaire Data

Descriptive statistics for the AQ and OLIFE can be seen in Table 1. Total autistic traits and total schizotypal traits were positively correlated (*r* = .305, *p* = .002, 95% CI = [.111, .499], BF10 = 12.03). The inter-trait sub-scale correlations of interest (Bonferroni *p* < .0025, Figure 2) revealed positive correlations between Social Skills and Introvertive Anhedonia (*r* = .528, *p* = 2.78e-08, 95% CI = [.355, .701], BF10 = 503158.12), between Cognitive Disorganization and both Attention Switching (*r* = .361, *p* = 2.78e-04, 95% CI = [.171, .551], BF10 = 85.14) and Communication (*r* = .383, *p* = 1.09e-04, 95% CI = [.195, .571], BF10 = 202.16), and between Communication and Impulsive Non-Conformity (*r* = .367, *p* = 2.16e-04, 95% CI = [.178, .557], BF10 = 107.35). There was a negative correlation between Imagination and Unusual Experiences (*r* = -.304, *p* = 2.46e-03, 95% CI = [-.110, -.498], BF10 = 11.62). Age did not correlate with scores on either the AQ (*r* = -.020, *p* = .843) or OLIFE (*r* = -.041, *p* = .690). AQ scores were higher in males (*m* = 18.4, *SD* = 5.7) than females (*m* = 15.2, *SD* = 5.4, *t*(95) = 2.86, *p* = .005). There were no sex differences in OLIFE scores (*t*(95) = .419, *p* = .676). The full correlation matrix can be seen in Supplementary Figure 1.

**Figure 2.**
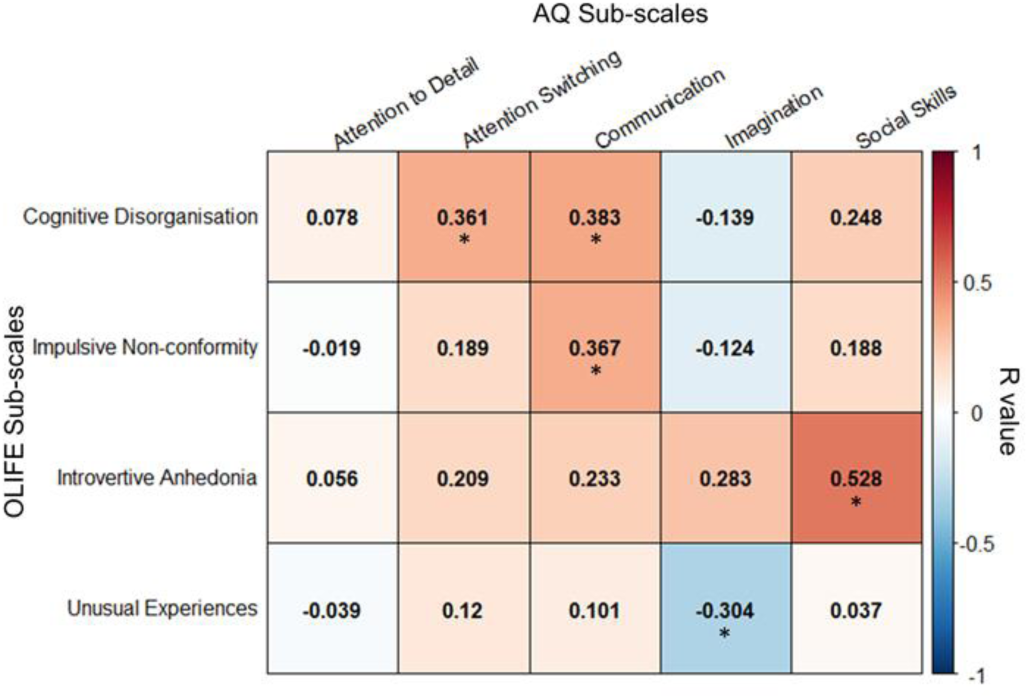
The inter-trait correlations between the sub-scales of the Autistic Spectrum Quotient and the Oxford-Liverpool Inventory of Feelings and Experiences. Correlations significant at Bonferroni corrected p < .0025 are marked with an asterisk.

**Table 1.**
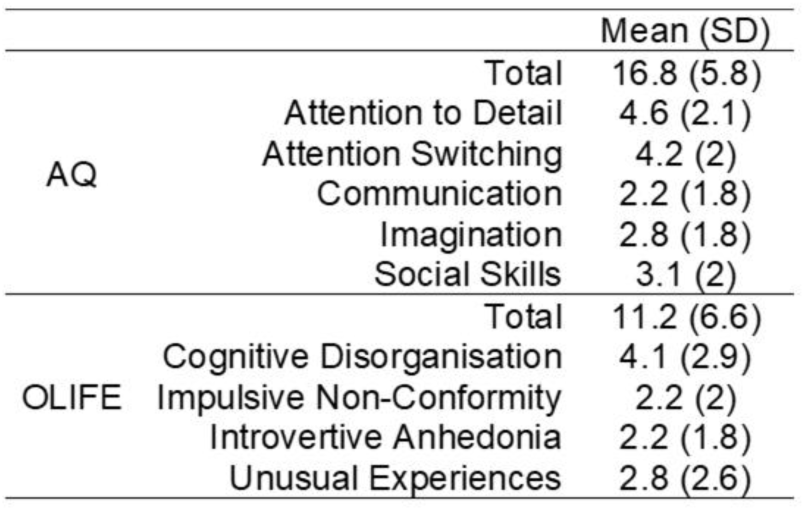
Autistic Spectrum Quotient and OLIFE scores in total and for each subscale

Overall, autistic traits and schizotypal traits were positively correlated, most importantly on the social skills and introversion sub-scales that may contribute to phenotypic convergence. These relationships were thus targeted for establishing similarities and differences in the neural activity and synchronization associated with biological motion perception. Exploratory analyses on the associations within the sub-scales of each respective trait questionnaire, and the inter-trait correlations are presented in the Supplementary Materials.

#### The relationship between neural activity and biological motion perception

Biological motion was associated with a widespread and distributed increase in neural activity (Figure 3A, FWE *p* < .001). Bilateral activation was evident in the lingual gyri, cuneus, precuneus, thalamus, precentral gyrus, superior, medial, and inferior frontal gyri, superior, middle, and inferior temporal gyri, fusiform gyri, lentiform nucleui. Unilateral activation was evident in the right superior and inferior frontal gyri. Cerebellar activation was observed in the bilateral cerebellar tonsil, unvula of vermis, and right declive. Negative relationships between biological motion and neural activity were found in bilateral caudate/parahippocampus, bilateral post-central gyri, right thalamus, and bilateral middle insula.

**Figure 3.**
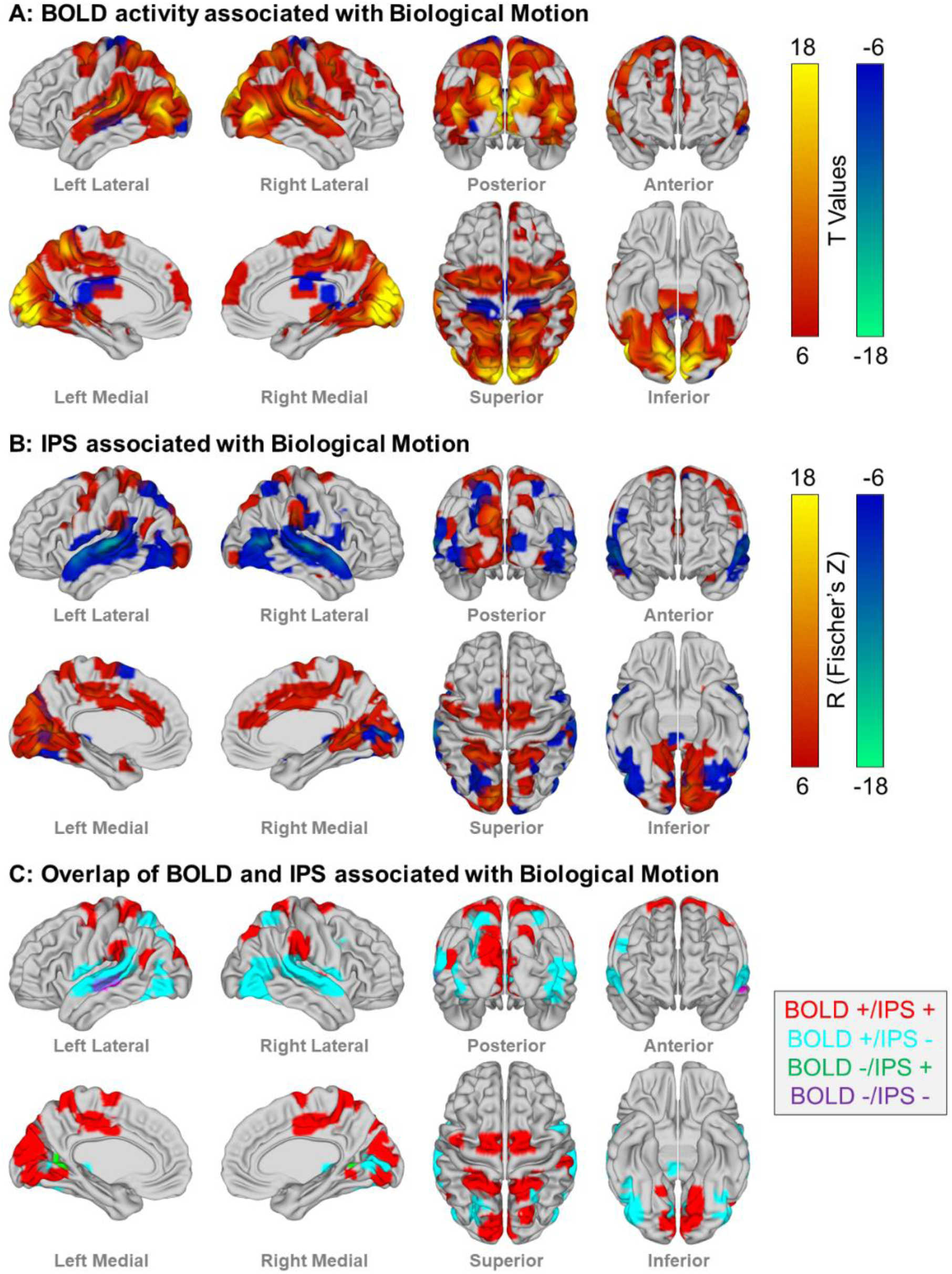
The convergent and divergent patterns of neural activity and synchronization associated with biological motion. A: Regions exhibiting an increase (red) or decrease (blue) in neural activity using GLM analyses with biological motion as a regressor (FWE *p* < .001). B: Regions exhibiting an increase (red) or decrease (blue) in inter-subject neural synchronization associated with biological motion (FWE *p* < .001). C: Logical overlays of regions exhibiting both neural activity and synchronization associated with biological motion, with relationships being convergent (positive or negative for both neural activity and synchronization) or divergent (positive and negative for either neural activity or synchronization).

#### The relationship between neural synchronization and biological motion perception

Biological motion was associated with an increase in inter-subject phase synchronization in a similarly distributed, but more discrete set of regions (Figure 3B, FWE *p* < .001) as was observed in the GLM analysis. Bilateral increases were observed in the lingual gyri, inferior parietal lobes, superior parietal lobes, middle temporal gyri, and anterior cingulate gyri.

Unilateral synchronization in the left hemisphere was observed in the cuneus, fusiform gyrus, precentral gyrus, superior frontal gyrus, and inferior frontal gyrus, and in the right hemisphere in the inferior occipital gyrus, precuneus, post-central gyrus, parahippocampus, posterior cingulate, middle cingulate gyrus, medial frontal gyrus, claustrum, uvulvua of vermis, and right declive. ISPS was negatively associated with biological motion bilaterally in the lingual gyri, fusiform gyri, cuneus, superior and middle temporal gyri, and superior parietal lobes. Right hemispheric decreases were evident in the pre and post-central gyri, inferior parietal lobe, thalamus, parahippocampus, and inferior frontal gyrus. Left hemispheric decreases were evident in the inferior and middle occipital gyri, inferior temporal gyrus, precuneus, posterior cingulate, and superior frontal gyrus.

### Overlapping neural activity and synchronization associated with biological motion perception

Logical overlays of the maps generated by the GLM and ISPS analyses reveal the convergent and divergent patterns of neural activity and synchronization associated with biological motion perception (Figure 3C).

#### GLM Positive/ISPS Positive

Both an increase in neural activity and ISPS was evident bilaterally in the precuneus, superior parietal lobe, precentral gyri, cuneus/lingual gyri, posterior cingulate gyrus, inferior parietal lobe, and left middle temporal gyrus and right claustrum.

#### GLM Positive/ISPS Negative

Several regions exhibited an increase in neural activity but a decrease in ISPS in response to biological motion, most notably in a large bilateral swathe along the superior temporal gyri, and also bilateral superior parietal lobe, bilateral fusiform gyrus, left cuneus and precuneus, right lingual gyrus, right middle temporal gyrus, and right inferior frontal gyrus.

#### GLM Negative/ISPS Positive

Only two small regions in bilateral parahippocampus exhibited a decreased neural activity and an increased ISPS associated with biological motion perception.

#### GLM Negative/ISPS Negative

A prominent region in the left middle temporal gyrus exhibited both a decreased neural activity and ISPS with biological motion perception, as did small regions in the left posterior cingulate and right parahippocampus.

### Autistic and Schizotypal traits associated with neural activity in response to biological motion

We next conducted a GLM analysis to establish the extent to which autistic and schizotypal traits are associated with neural activity in response to biological motion (Figure 4A). The first level analyses with biological motion as a regressor were entered into a second-level analysis with trait scores as a regressor. Autistic traits, with schizotypal traits as a covariate, were negatively correlated with the neural response to biological motion in a cluster (k = 187) with two peak voxels in the left and right middle cingulate gyrus. The AQ sub-scale of Social Skills, with the OLIFE sub-scale of Introversion entered as a covariate, was negatively associated with the neural response to biological motion in a cluster (k = 157) with two peak voxels in the right precuneus. The converse analyses showed that neural activity associated with biological motion perception was not associated with schizotypal traits (with autistic traits as a covariate), nor with the Introversion sub-scale (with the Social Skills sub-scale as a covariate).

**Figure 4.**
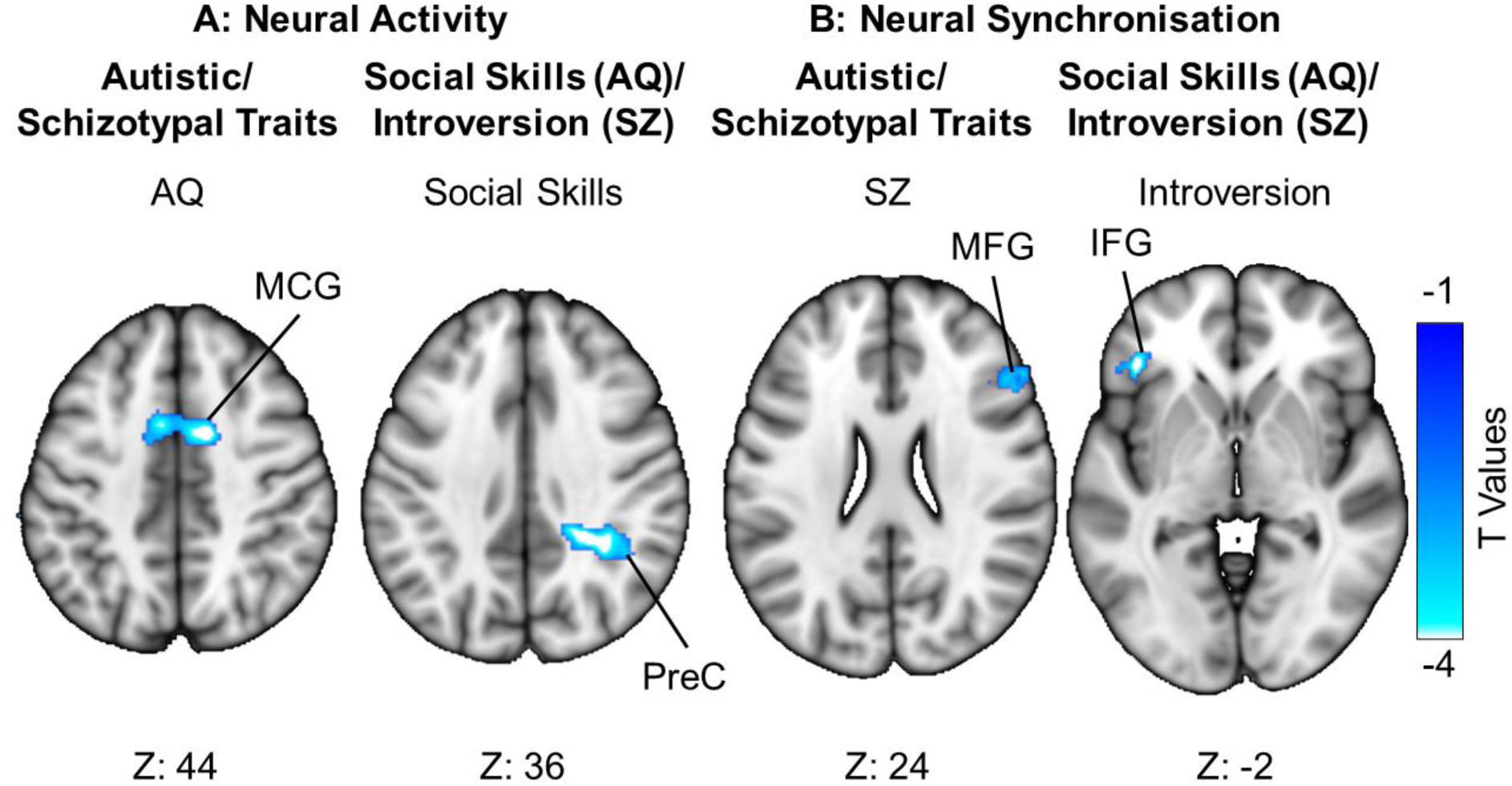
Patterns of neural activity and synchronization associated with biological motion dissociate autistic and schizotypal traits. A: The neural activity associated with biological motion decreased with increasing autistic traits (with schizotypal traits as an orthogonal regressor) in the middle cingulate gyrus (MCG), and decreased with increasing atypical social skills (with introversion as an orthogonal regressor) in the Precuneus (PreC). B: Neural synchronization associated with biological motion decreased with increasing schizotypal traits (with autistic traits as an orthogonal regressor) in the middle frontal gyrus (MFG) and decreased with increasing introversion (with social skills as an orthogonal regressor) in the inferior frontal gyrus (IFG). For visualization purposes, these figures depict an uncorrected voxel threshold of *p* < .01, followed by a FWE cluster threshold of *p* < .05.

### Autistic and Schizotypal traits associated with neural synchronization in response to biological motion

The relationship between ISPS and biological motion decreased with schizotypal traits, with autistic traits as a covariate, in a cluster (k = 39) with two peak voxels in the right middle frontal gyrus. The OLIFE subscale of Introversion, with the AQ sub-scale of Social Skills as a covariate, was negatively associated with the relationship between ISPS and biological motion in a cluster (k = 39) with a peak voxel in the left inferior frontal gyrus (Figure 4B). The converse analyses showed that neural synchronization associated with biological motion perception was not associated with autistic traits (with schizotypal traits as a covariate), nor with the Social Skills sub-scale (with the Introversion sub-scale as a covariate)

#### Individual differences in autistic and schizotypal traits in neural synchronization

Descriptive statistics for the extent of inter-subject phase synchronization of neural activity across the sample, independent of stimulus features, are depicted in Figure 5. Neural synchronization was greatest in posterior visual areas and auditory regions in the temporal cortex, as would be expected given that all participants are viewing and listening to the same stimulus.

**Figure 5.**
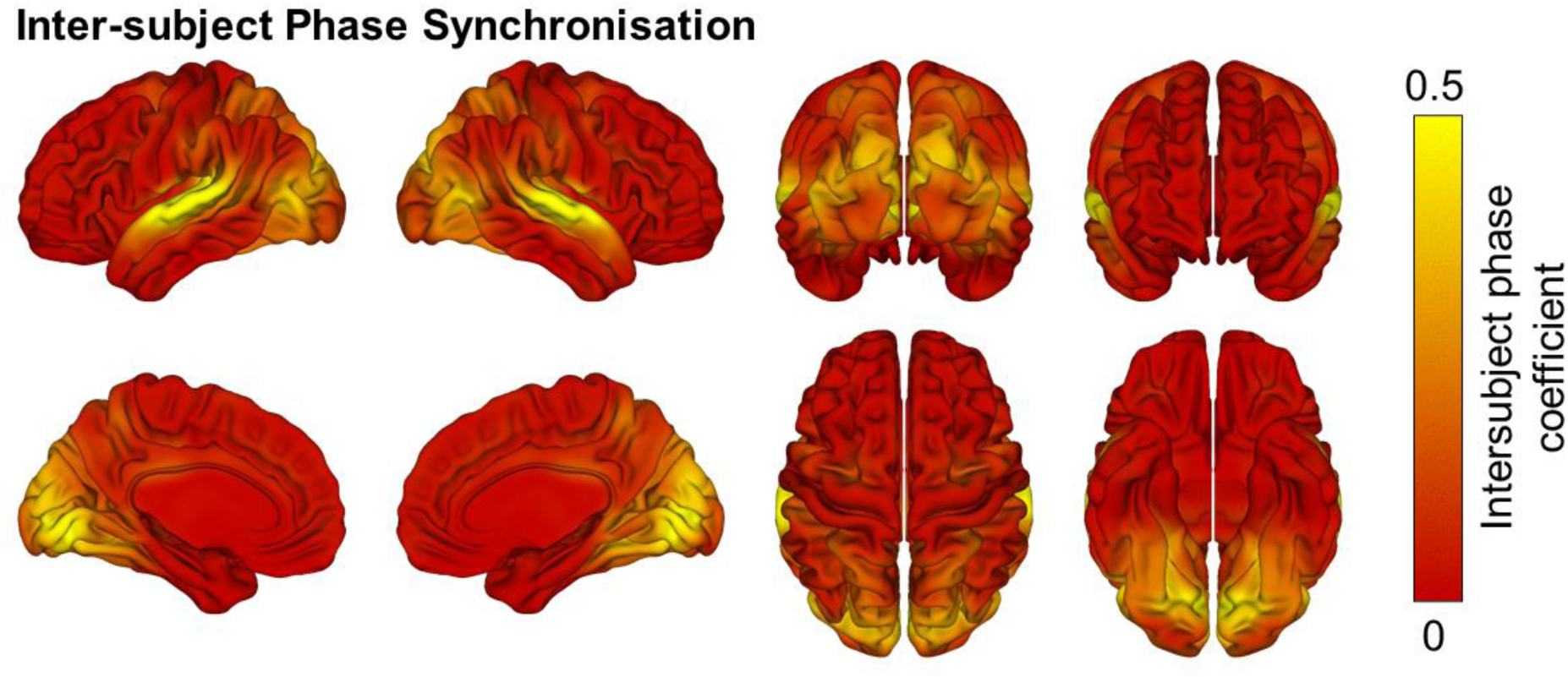
Inter-subject phase synchronization of neural activity across the sample, independent of stimulus features. Of interest is how this neural synchronization varies with autistic and schizotypal traits in general, and social skills and introversion specifically, across the whole stimulus run, irrespective of biological motion content. Autistic and schizotypal traits, which are positively correlated, exhibit divergent patterns of ISPS (Figure 6), with autistic traits positively correlating with ISPS in a cluster (k = 86) with three peak voxels in the left superior temporal gyrus, and increasing schizotypal traits associated with a decrease in ISPS in a cluster (k = 64) with three peak voxels in the right precuneus. Despite a positive correlation between the Social Skills sub-scale of the AQ and the Introversion sub-scale of the OLIFE, neither were associated with ISPS when entered as regressors.

**Figure 6.**
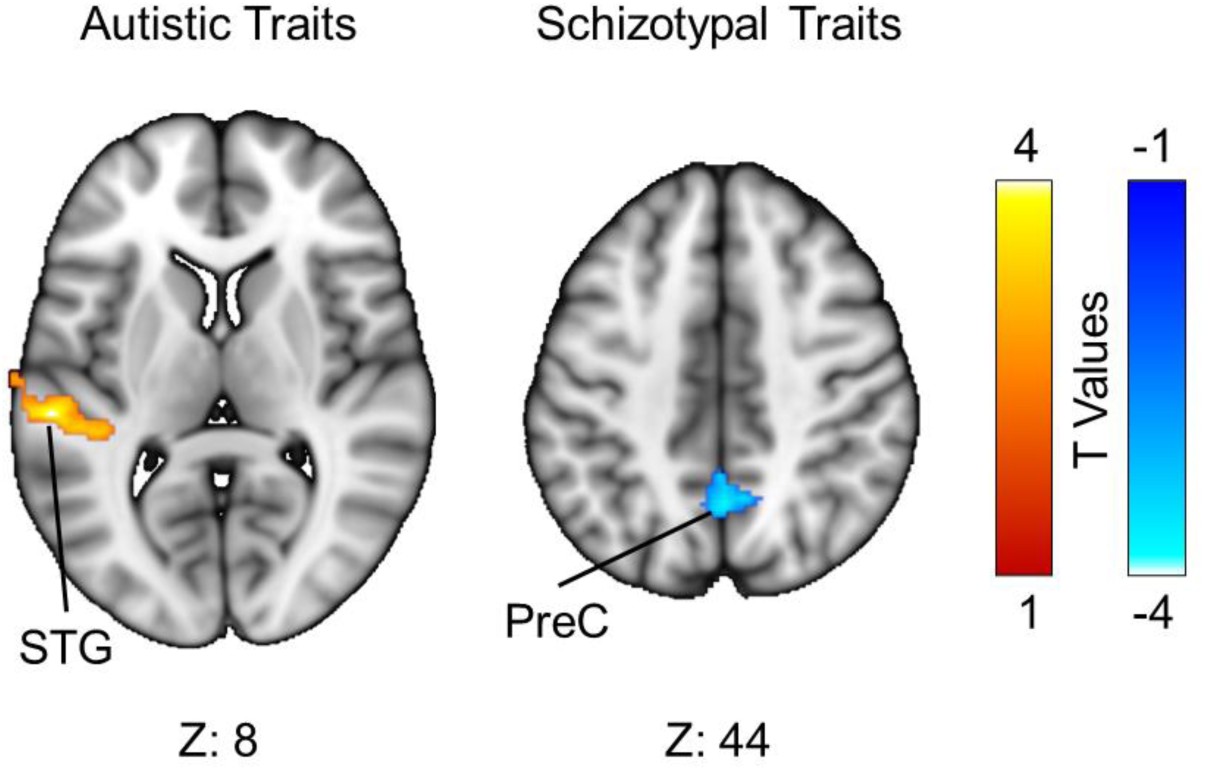
The relationship between overall autistic and schizotypal traits and inter-subject phase synchronization independently of stimulus features. Autistic and schizotypal traits were entered as orthogonal regressors against the degree of inter-subject phase synchronization, across the stimulus run. Neural synchronization increased with increasing autistic traits in the Superior Temporal Gyrus (STG), and decreased with increasing schizotypal traits in the Precuneus (PreC). For visualization purposes, these figures depict an uncorrected voxel threshold of *p* < .01, followed by a FWE cluster threshold of *p* < .05.

### Functional connectivity (seed-based phase synchronization) associated with biological motion perception

At a strict threshold of FDR *p* < .001, there was a significant negative association between biological motion and the synchronization between the right fusiform gyrus and right lateral occipital cortex. That is, as biological motion content increased, neural activity in these regions became desynchronized. A more liberal threshold of FDR *p* < .01 revealed further relationships (Figure 7). The right lateral occipital cortex showed an additional decrease in synchronization with the left fusiform gyrus as biological motion increased. The right superior parietal lobe exhibited decreased neural synchronization with both the right operculum inferior frontal gyrus and right insula. There was a decrease of neural synchronization between the right lateral orbital gyrus and right ventral inferior frontal gyrus, between the right inferior temporal gyrus and right paracentral lobule, and between the right middle frontal gyrus and left postcentral gyrus. Lastly, there was an increase in neural synchronization between the right medial orbital gyrus and left superior parietal lobe as biological motion content increased.

**Figure 7.**
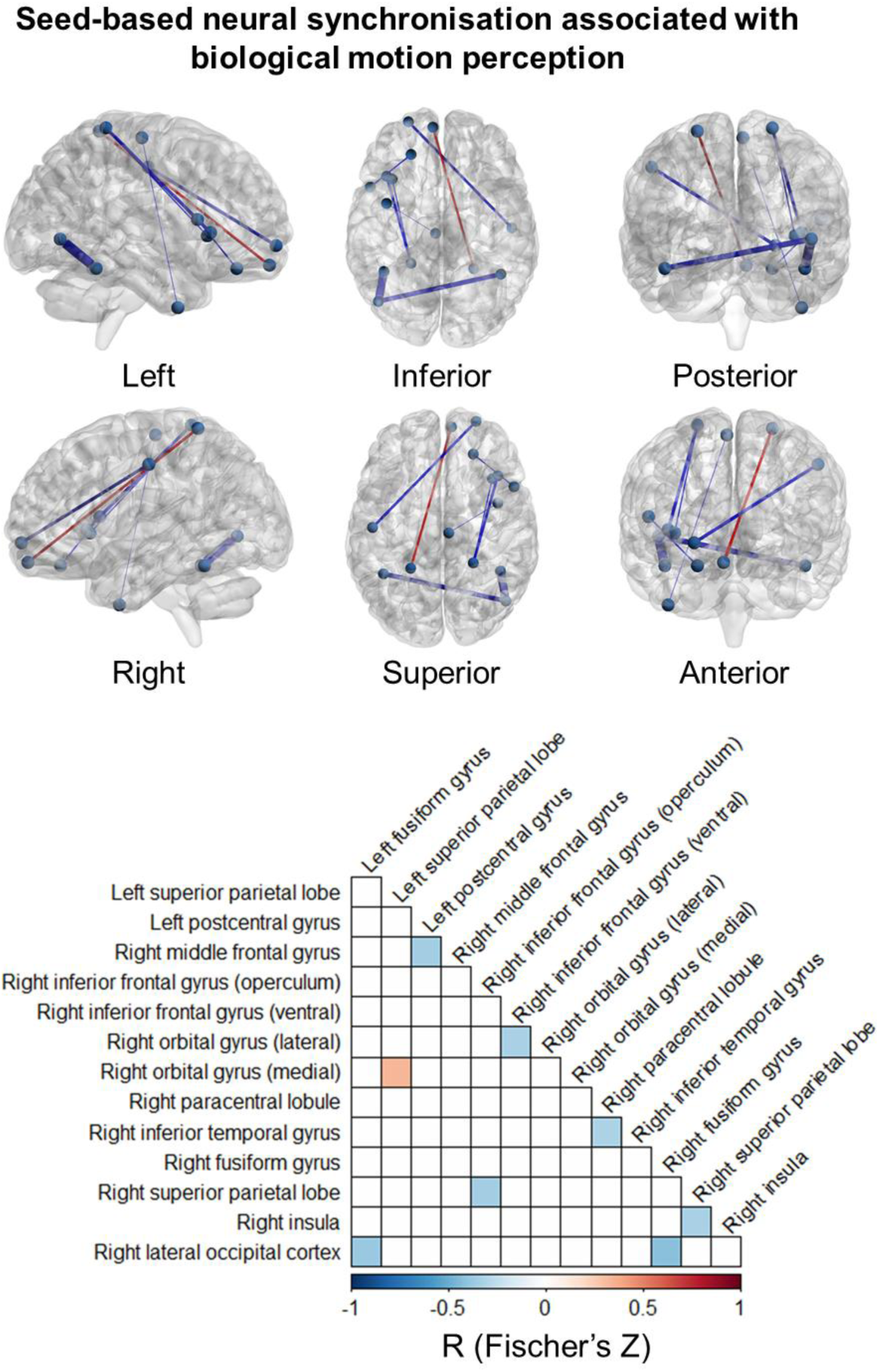
Functional connectivity associated with biological motion perception (FDR *p* <.01). Phase synchronization of neural activity between the right medial Orbital Gyrus and left Superior Parietal Lobe increased with biological motion, whilst all other region pairs exhibited a decreased neural synchronization, including between the right fusiform gyrus and right lateral occipital cortex (FDR *p* <.001). The relationship between neural synchronization between regions and biological motion was not associated with individual differences in autistic and schizotypal traits

Additional analyses investigated whether seed-based phase synchronization across the whole stimulus run, irrespective of biological motion content, was associated with individual differences in autistic and schizotypal traits, but no results survived correction for multiple comparisons. Likewise, there was no evidence that the relationship between neural synchronization between brain regions and biological motion content varied with autistic and schizotypal traits when correcting for multiple comparisons.

## Discussion

Our main finding was that, whilst neural activity associated with the perception of biological motion increases in a widespread network of brain regions, the synchronization of neural activity between individuals differs within this network. Biological motion perception was associated with increased neural synchronization in visual and parietal regions, whereas de-synchronization was evident in temporal and frontal regions. We further found that autistic traits are associated with changes in overall neural activity related to biological motion in the precuneus and middle cingulate gyrus, whilst schizotypal traits are associated with changes in neural synchronization in the middle and inferior frontal gyri. These results suggest that correlated autistic and schizotypal traits, especially those relating to social behavior, are associated with different neural responses to social stimuli, namely the magnitude of neural activity for autistic traits, and the between-individual synchronization of neural activity for schizotypal traits.

### Convergent and divergent patterns of neural activity and synchronization in relation to biological motion perception

The perception of biological motion was associated with an increase in neural activity in an extensive network of regions encompassing the occipital, temporal, parietal and frontal cortices. This distributed pattern of neural activity characterizes the well-established ‘social brain’ implicated in action observation (occipital cortex, posterior temporal regions, parietal, and pre-motor regions) and mentalizing (anterior-temporal, temporal-parietal junction, medial-frontal gyri) (Adolphs, 2009; Li et al., 2018). However, the distribution of neural synchronization between individuals during biological motion perception varied within this network. Regions in primary visual areas, face and body selective visual areas, inferior parietal lobe, pre-central gyri, temporal-parietal junction, and cingulate cortex showed an increase in neural activity that is also highly synchronized across individuals. In contrast, the superior parietal lobe, superior and middle temporal gyri, fusiform gyri, and inferior frontal gyri exhibited de-synchronization of neural activity, despite an overall increase in neural activity.

Broadly speaking, neural synchronization varied along a posterior – anterior axis, with increased synchronization observed in posterior visual areas and the parietal lobe, whereas decreased synchronization was observed in temporal association regions and frontal regions. This agrees with previous findings of a systematic gradient of neural reliability (Kauppi, Jääskeläinen, Sams, & Tohka, 2010; Kauppi, Pajula, Niemi, Hari, & Tohka, 2017) that reflects a global cortical hierarchy of parsing, integration, and prediction of information at different timescales (Baldassano, Chen, Zadbood, Pillow, Hasson, & Norman, 2017; Hasson, Chen, & Honey, 2015; Hasson, Malach, & Heeger, 2010; Hasson, Yang, Vallines, Heeger, & Rubin, 2008; Huntenburg, Bazin, & Margulies, 2018; Kiebel, Daunizeau, & Friston, 2008). Sensorimotor areas operate at short timescales and reflect stimulus features, therefore showing a high degree of neural reliability when participants are viewing the same stimulus. Neural activity is less reliable in temporal and prefrontal regions, which are involved in idiosyncratic interpretations of visual input and decisions about how to act, and which operate at longer timescales. Disturbances in the extent of this gradient have also been observed in autism (Watanabe, Rees, & Masuda, 2019) and psychosis (Wengler, Goldberg, Chahine, & Horga, 2020).

This dichotomy may provide a key insight into the neural mechanisms underpinning biological motion perception. The perception of biological motion in visual and parietal areas is tightly coupled to sensory stimulus features, which is shared across all participants. These include visible body parts, the extent of motion, or the specific action, and how these change on a moment-to-moment basis, and their perception leads to consistent neural responses across subjects. In contrast, neural activity in temporal and frontal regions became increasingly less reliable during the perception of biological motion (despite overall increase in activity in the BOLD-GLM analysis), possibly reflecting more subjective and idiosyncratic interpretations about the other’s behavior. This dissociation includes the IPL and the IFG, which are implicated in visuomotor transformations of observed actions (the mirror-matching system, Molenberghs, Cunnington, & Mattingley, 2012), and which exhibited increased neural activity during biological motion perception. This division is consistent with previous models of mirror neuron function, either retrieving associated goals from observed actions (Rizzolatti & Craighero, 2004), or predicting future actions from inferred goals (Kilner, Friston, & Frith, 2007), wherein the motor response of the IPL is associated with stimulus features such as the low-level kinematic features of the observed action, whereas that of the IFG is associated with the interpreted goals of the action.

These results highlight that analysis of BOLD response amplitudes as well as inter-subject similarity of the responses provide complimentary information on the neural mechanisms of the perception of social stimuli. This is further evident in the observation that several regions that exhibited biological motion related synchronization did not exhibit concurrent changes in overall neural activity, and would not otherwise have been associated with biological motion. These included increased synchronization in extensive bilateral regions encompassing the lingual gyri and parahippocampus, along with small regions in anterior cingulate and inferior occipital gyri, which may be involved in the perception of stimulus features that are not necessarily social in nature, such as motion itself. Decreased synchronization in the absence of a change in overall neural activity was evident bilaterally along the entire superior temporal gyri and sulci, as well as small bilateral regions in the fusiform gyri, which may be involved in individual high-level interpretations which follow biological motion perception, but which are not directly related to it, such as the overall intentions of the individual or the context of the interaction.

The analysis of phase synchronization of neural activity also revealed new insights into the functional connectivity of these networks during biological motion perception. The network of regions involved in social functions involves several nested networks working in parallel to perform different roles, such as mentalizing, face perception, and empathy (Li et al., 2018; Sokolov, Zeidman, Erb, Ryvlin, Friston, & Pavlova, 2018). However, we found that the synchronization of neural activity between regions decreased during biological motion perception. Therefore, whilst the functional significance of each region is related to the extent of biological motion, and to the input of other regions, there is a greater reduction in instantaneous neural reliability. Phase de-synchronization of neural activity between regions has been established as a result of inhibitory coupling or delayed relay of information (Li & Zhou, 2011, Protachevicz, Hansen, Iarosz, Caldas, Batista, & Kurths, 2021). The parallel functions operating within the social brain network may elicit greater differentiation of neural activity between regions as biological motion perception increases, thus causing the apparent neural de-synchronization.

### Neural activity and synchronization differentiate individual differences in autistic and schizotypal traits

Autistic and schizotypal traits were positively correlated, especially on measures of social behavior, but also on more general indices of cognitive organization and control. Only traits relating to imagination and unusual experiences (homologous to positive schizotypal traits) were negatively correlated. These findings support previous theoretical and empirical work suggesting an intersection of features of autism and schizophrenia, and related traits in the neurotypical population. However, it is not clear if these traits share a common mechanism. The divergent pattern of autism and schizotype-dependent neural activity and synchronization in response to biological motion perception suggests that these traits do reflect phenotypic overlap with separate neural bases.

The increased synchronization with autistic traits in the superior temporal gyrus was observed by Salmi et al., (2009), and the decreased synchronization in the precuneus with increasing schizophrenic symptoms was observed by Lerner et al., (2018). It is unclear exactly how these differential patterns of neural synchronization contribute to autistic and schizotypal traits, however, the specificity of these replications warrants further investigation.

Analyzing the relationship between neural response amplitudes versus synchronization driven by biological motion perception yielded a clear dissociation between autistic and schizotypal traits. Autistic traits, and social skills in particular, were associated with deceases in overall neural activity related to biological motion, whereas schizotypal traits, and introversion in particular, were associated with decreases in neural synchronization. The social difficulties associated with autism are in part reflected in reduced overall neural activity in response to the perception of complex social emotional interactions, which may reflect or cause an insensitivity to such stimuli compared to those with schizophrenia (Sasson et al., 2007). However, whilst watching those same interactions, the social difficulties associated with schizophrenia correspond with decreased neural synchronization between individuals, which may reflect not only a reduction in short-term integration of stimulus features, but also an inability to establish neural ‘rapport’ with others that enables mutual psychological states, and which may contribute to spurious mental state attributions (Ciaramidaro et al., 2015). Moreover, the decreased synchronization associated with schizotypal traits was focused in frontal areas (the middle and inferior frontal gyri), which are associated in schizophrenia with high-level inferences and decision making in social interactions (Li, Chan, Gong, Liu, Liu, Shum, & Ma, 2012; Russell, Rubia, Bullmore, Soni, Suckling, Brammer, Simmons, Williams, & Sharma, 2000; Shin, Choi, Lee, Shin, Jang, & Kim, 2015; Takei, Suda, Aoyama, Yamaguchi, Sakurai, Narita, Fukuda, & Mikuni, 2013). In contrast, decreased neural activity associated with autistic traits was observed in middle cingulate gyrus and precuneus, both of which have been implicated in autism with a reduced awareness of one’s own actions and decisions in social interactions and distinguishing self from others (Chiu, Kayali, Kishida, Tomlin, Klinger, Klinger, & Montague, 2008; Just, Cherkassky, Buchweitz, Keller, & Mitchell, 2014; Lombardo, Chakrabarti, Bullmore, Sadek, Pasco, Wheelwright, Suckling, MRC AIMS Consortium, Baron-Cohen, 2010; Martineau, Andersson, Barthélémy, Cottier, & Destrieux, 2010; Tomlinm, Kayali, King-Casascedric, Camerer, Quartz, & Montague, 2006). These regions were not associated with the perception of biological motion at the population level (see also Puglia & Morris, 2017), suggesting that social difficulties in autism and schizophrenia may result from downstream or upstream secondary processes that rely on, but are not directly implicated in, the perception of others behavior.

### Limitations

Self-report survey measures of autistic and schizotypal traits in the neurotypical population are consistent with the with spectrum models of autism and psychosis, but assume that these traits are conceptually aligned with these conditions in a linear and unidimensional distribution across the spectrums (Sasson & Bottema-Beutel, 2021). Autism and psychosis-spectrum disorders are multi-dimensional and exhibit qualitative differences to the traits measured in these surveys, and so the extent to which the current results can be extrapolated either quantitatively or qualitatively to those with autism or schizophrenia remains to be seen. Moreover, autistic people find social interactions to be easier when they are interacting with other autistic people (Bolis, Lahnakoski, Seidel, Tamm, & Schilbach, 2020; Crompton, Hallett, Ropar, Flynn, & Fletcher -Watson, 2020), ostensibly due to an increased compatibility in the styles of thinking and behavior that make social interactions more fluid and empathetic (Bolis, Balsters, Wenderoth, Becchio, & Schilbach, 2017; Milton, 2012). In the current study, the behavior of neuroptypical people were employed as stimuli, and so further research should examine whether the neural response to the behavior of others is affected by the similarities of neurophenotypes between individuals.

Furthermore, although we aimed to reflect naturalistic conditions by having participants freely and passively view a complex and dynamic social stimulus, it is not possible to interpret the functional significance of neural (de)synchronization associated with biological motion perception, nor the activity of regions in which the neural profile was associated with schizotypal and autistic traits. These regions have been previously implicated in socio-cognitive performance in autism and schizophrenia, but how their differing neural profiles contribute to convergent behavioral phenotypes requires controlled experimental manipulation. Moreover, as eye movements were not measured, it is not possible to assess the differing attention resources allocated to the stimulus, and how these may be associated with autistic and schizotypal traits (although attentional and cognitive sub-scales of these trait measures did not correlate with those relating to social behavior, and were associated with different neural responses, as reported in the supplementary material).

### Conclusion

The perception of biological motion elicits both overlapping and dissociable patterns of neural activity and neural synchronization between individuals. Increases in neural synchronization were observed primarily in regions of the brain associated with stimulus processing (visual, motor regions), whereas decreases in neural synchronization were observed primarily in regions associated with interpretation and decision making (temporal and frontal regions). These differences correspond to a well-established posterior-anterior axis of neural reliability implicated in temporal parsing, integration, and prediction, that we can now apply to the perception of social interactions. Moreover, patterns of neural activity and synchronization differentiated the highly correlated individual differences in autistic and schizotypal traits of the large sample of the neurotypical population in regions that were not directly implicated in biological perception, but which have previously been implicated in social functions. The highly convergent individual differences in social behavior that correspond to autistic and schizotypal traits, and by possible extension the common social difficulties encountered by those with autism and schizophrenia themselves, do not reflect a shared etiology, but disparate mechanisms that elicit superficially similar phenotypes. The use of complex and naturalistic social interactions provides new avenues for future research. Different temporal profiles of neural activity can be dissociated by the same stimulus properties, in this case the perception of other people’s behavior, and this can reveal different neural mechanisms associated with autistic and schizotypal individual differences that cannot be distinguished at the behavioral level.

## Supplementary Materials

**Supplementary Figure 1.**
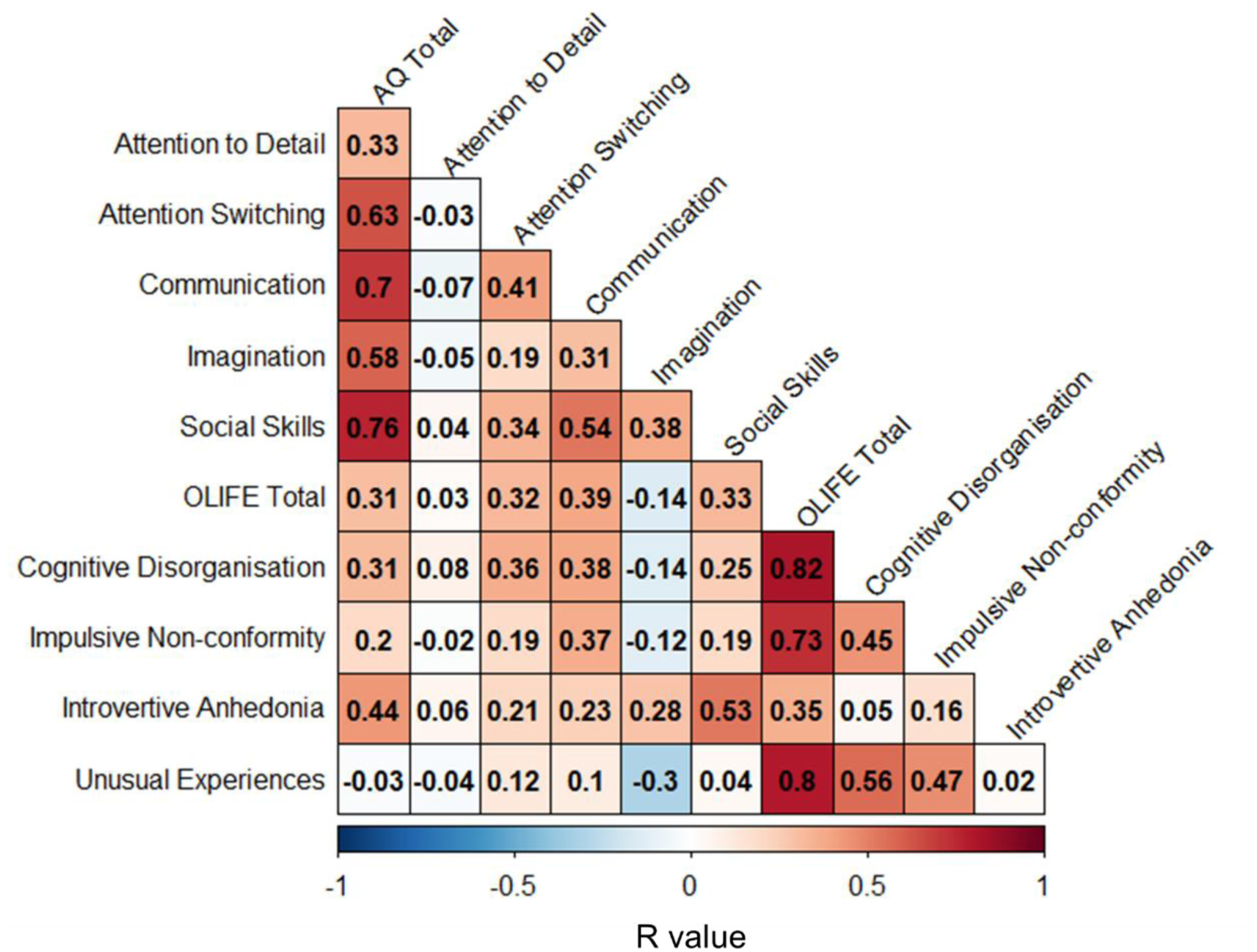
Correlation matrix showing the within trait and between trait relationships between overall autistic and schizotypal traits and the respective sub-scales.

## Supplementary Analysis

The main hypotheses of this study predicted that there would be a positive corelation between autistic and schizotypal traits, especially with respect to those subscales measuring social behaviours, and that despite these positive relationships, the differential pattern of neural synchronisation and activity may dissociate these convergent traits. This is indeed what we report. We also found several other facets of autistic and schizotypal traits that showed inter-trait correlations. These were positive relationships between Cognitive Disorganisation, Attention Switching, and Communication, between Communication and Impulsive Non-Conformity, and a negative relationship between Imagination and Unusual Experiences., but Although these did not measure social behaviours, we nevertheless conducted exploratory analyses to assess the degree to which these correlated traits may also be differentiated by the neural activity and synchronisation associated with biological motion perception. In addition, we also investigated whether the neural activity and synchronisation associated with biological motion perception could distinguish intra-trait relationships between the subscales, by entering all subscales for each trait measure in separate analyses.

We first examined intra and inter-trait relationships with neural synchronisation overall, irrespective of stimulus features, followed by examining intra and inter-trait relationships with neural activity associated with biological motion perception, and lastly, the intra and inter-trait relationships with neural synchronisation associated with biological motion perception.

### Individual differences in autistic and schizoptyal traits in neural synchronisation

#### Intra-Trait analyses (p < 001, cluster corrected)

The five sub-scales of the Autistic Spectrum Quotient were entered as regressors to establish if they were associated with the degree of IPS across the whole stimulus run, irrespective of biological motion content.

*Autistic Traits (Supp Figure 2A):* Attention Switching was positively correlated with IPS in one cluster (k = 241) with peak voxels in the right posterior insula cortex and claustrum. IPS decreased with increasing Social Skills deficits in one cluster (k = 73) in the inferior semi-lunar lobe of the cerebellum.

*Schizotypal Traits (Supp Figure 2B):* The four sub-scales of the Oxford-Liverpool Inventory of Feelings and Experiences were entered as regressors to establish how individual differences in schizotypal traits are associated with IPS. Increasing traits of Impulsive Non-conformity were associated with decreasing IPS in one cluster (k = 100) with a peak voxel in the right precuneus. A decrease in IPS was also associated with increasing Unusual Experiences in two clusters (k = 72) with two peak voxels in the left cuneus, and three in the right precuneus.

#### Inter-Trait analyses (p < 001, cluster corrected)

The AQ sub-scale of Communication and the OLIFE sub-scale of Impulsive Non-conformity, whilst positively correlated, revealed different patterns of IPS (Supp Figure 2C). Communication difficulties were associated with increased IPS in two clusters (k = 70) with peak voxels in the right sub-gyral and left superior temporal gyrus, whereas increasing Impulsive non-conformity was associated with decreased IPS in a cluster (k = 108) with peak voxels in bilateral precuneus and left middle temporal gyrus.

The negative relationship between the Imagination sub-scale of the AQ and Unusual Experiences sub-scale of the OLIFE was evident in different patterns of IPS (Supp Figure 2D), with increasing difficulties in imagination being associated with increased IPS in a cluster (k = 73) with two peak voxels in the right middle frontal gyrus, and higher Unusual Experiences associated with decreased IPS in a cluster (k = 100) with two peak voxels in the right precuneus.

Entering the AQ sub-scales of Attention Switching and Communication and the OLIFE sub-scale of Cognitive Disorganisation as regressors revealed an increased IPS associated with Attention Switching in two clusters (k = 166), the first of which with peak voxels in the right posterior insula and superior temporal gyrus, the second of which with peak voxels in the left superior temporal gyrus and transverse temporal gyrus (Supp Figure 2E).

**Supplementary Figure 2.**
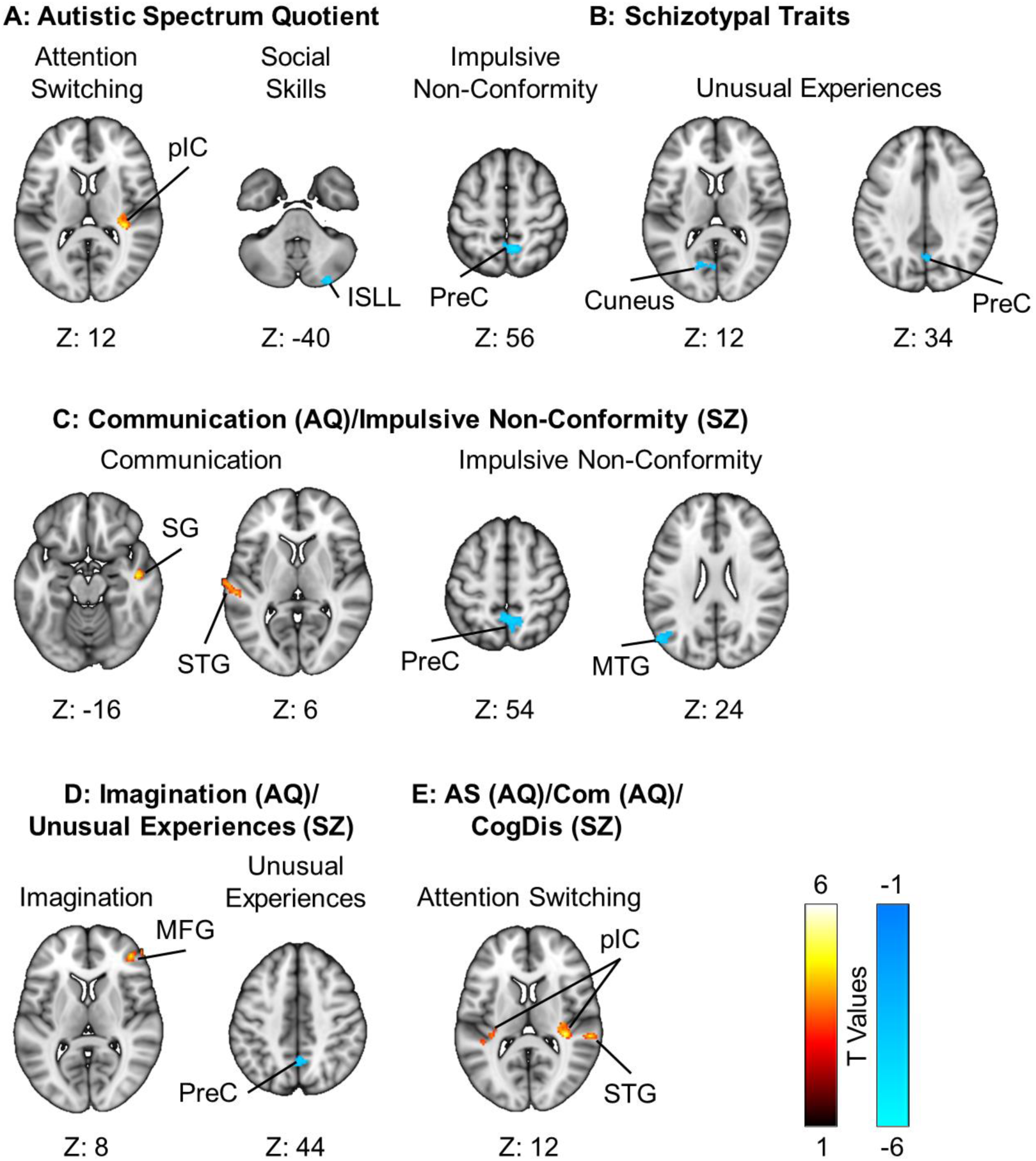
Inter-subject phase synchronisation of neural activity is related to individual differences in autistic traits (A) and schizotypal traits (B), and can differentiate inter-trait relationships between Communication and Impulsive Non-Conformity (C), Imagination and Unusual Experience (D), and Attention Switching, Communication, and Cognitive Disorganisation (E). Abbreviations: pIC = posterior insula cortex, ISLL = inferior semi-lunar lobe, PreC = Precuneus, SG = sub-gyral, STG = superior temporal gyrus, MTG = middle temporal gyrus, MFG = middle frontal gyrus.

### Autistic and Schizotypal traits associated with neural activity in response to biological motion

We next conducted a GLM analysis to establish the extent to which autistic and schizoptyal traits are associated with neural activity in response to biological motion. The first level analysis with biological motion as a regressor were entered into a second-level analysis with trait scores as a regressor.

#### Intra-Trait analyses (p < 001, cluster corrected)

When all five sub-scales of the AQ were entered as regressors, the sub-scale of Imagination showed a negative relationship with the neural response to biological motion in a cluster (k = 157) with three peak voxels in the left lingual gyrus. When all four sub-scales of the OLIFE were entered, the sub-scale of Impulsive Non-Conformity showed a positive relationship with the neural response to biological motion in a cluster (k = 110) with a peak voxel in the left post-central gyrus.

#### Inter-Trait analyses (p < 001, cluster corrected)

The OLIFE sub-scale of Impulsive Non-Conformity, with the AQ sub-scale of Communication as a covariate, was positively associated with the neural response to biological motion in a cluster (k = 111) with two peak voxels in the left post-central gyrus.

**Supplementary Figure 3.**
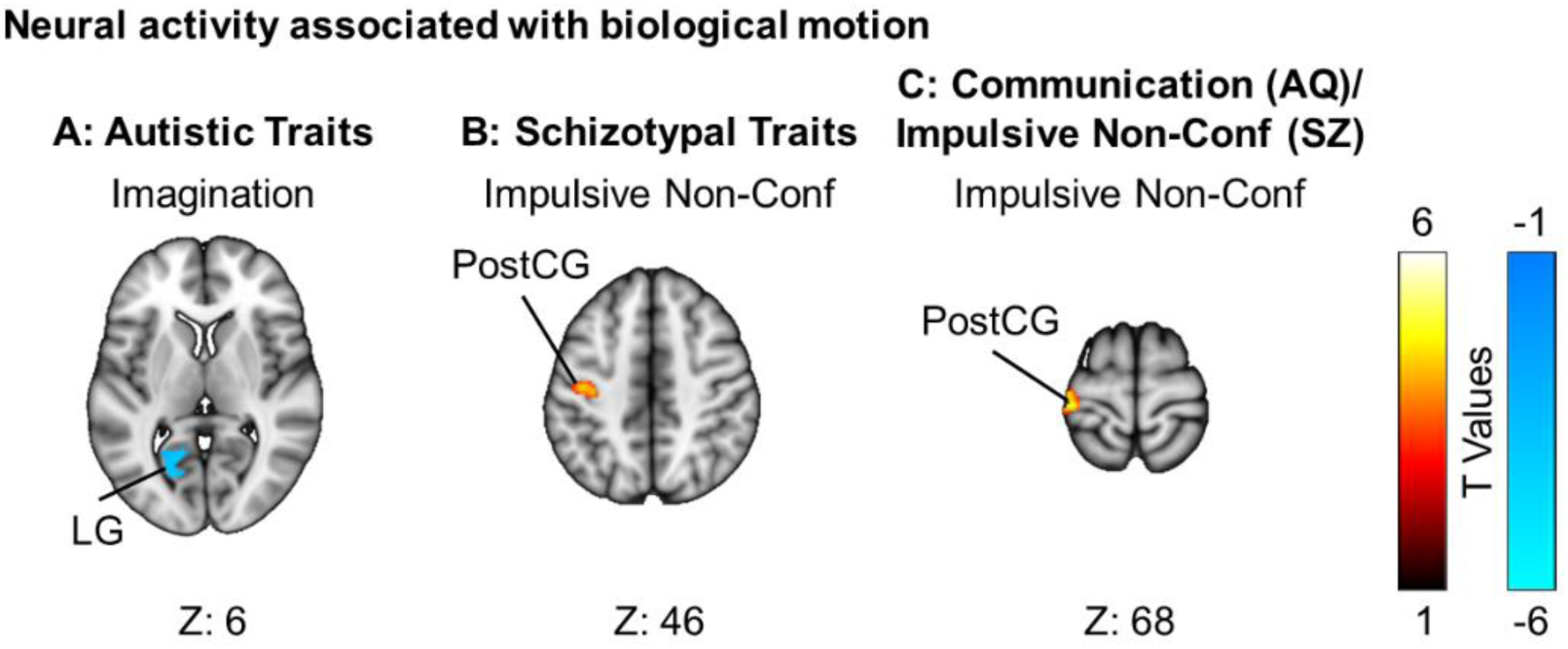
The magnitude of neural activity associated with biological motion perception varies with individual differences in autistic traits of imagination when other autistic traits are controlled for (A), and the schizotypal trait of impulsive non-conformity when other schizotypal traits are controlled for (B) and when the correlated autistic trait of communication is controlled for (C). Abbreviations: LG = lingual gyrus, PostCG = post central gyrus

### Autistic and Schizotypal traits associated with neural synchronisation in response to biological motion

For each individual the inter-subject phase synchronisation was correlated with biological motion for each voxel, and the r values were Fischer z transformed. These first-level results were entered into a second level analyses with trait scores as regressors.

#### Intra-Trait analyses (p < 001, cluster corrected)

With all AQ sub-scales entered as regressors, the relationship between IPS and biological motion increased with Imagination scores in a cluster (k = 49) with a peak voxel in the right superior parietal lobe, and with Social Skills scores in a cluster (k = 71) with three peak voxels in the right superior and middle temporal gyri. With all OLIFE sub-scales entered as regressors, the relationship between IPS and biological motion increased with Introversion scores in a cluster (k = 70) with three peak voxels in the right superior and medial frontal gyri.

#### Inter-Trait analysis (p < 001, cluster corrected)

The relationship between IPS and biological motion increased with scores on the AQ sub-scale of Communication, with the OLIFE sub-scale of Impulsive Non-Conformity, in a cluster (k = 38) in a cluster with a peak voxel in the left posterior insula cortex. The relationship between IPS and biological motion increased with the AQ sub-scale of Imagination, with the OLIFE sub-scale of Unusual Experiences as a covariate, in a cluster (k = 40) with a peak voxel in the right superior parietal lobe.

**Supplementary Figure 4.**
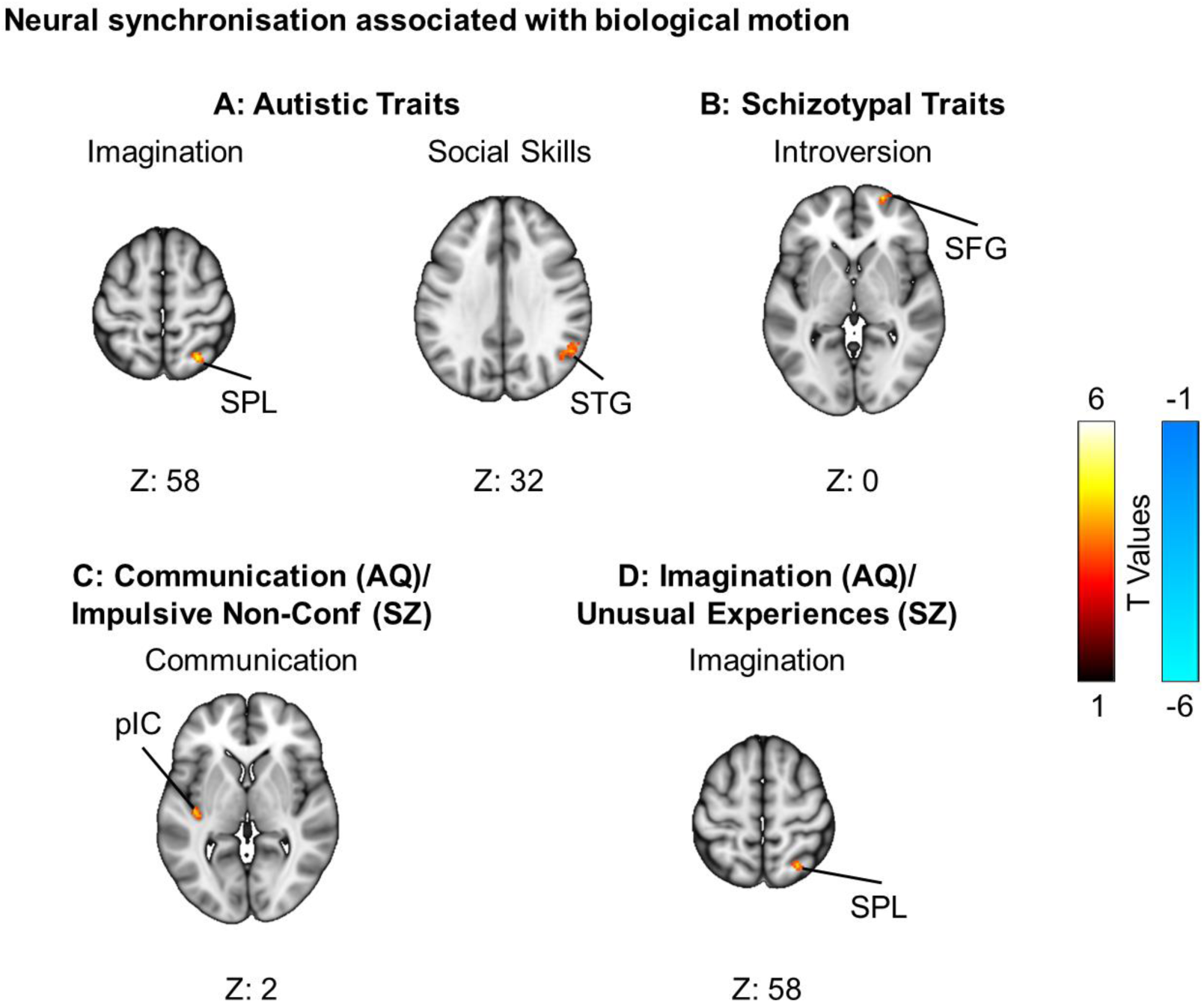
Neural synchronisation associated with biological motion perception varies with individual differences in autistic traits and schizotypal traits. The autistic traits of imagination and social skills are associated with increased neural synchronisation when other autistic traits are controlled for (A), and the schizotypal trait of introversion is associated with increase neural synchronisation when other schizotypal traits are controlled for (B). Neural synchronisation increased with the autistic traits of communication (when the correlated schizotypal trait of impulsive non-conformity was controlled) (C), and imagination (when the correlated schizotypal trait of unusual experiences was controlled for) (D). Abbreviations: SPL = superior parietal lobe, STG = superior temporal gyrus, SFG = superior frontal gyrus, pIC = posteior insular cortex.

